# Umbilical cord occlusions in near-term ovine fetus induce increased beat-to-beat heart rate variability correlating to decreases in neuroinflammation: a case for the afferent cholinergic anti-inflammatory pathway?

**DOI:** 10.1101/013169

**Authors:** M. G. Frasch, M. Szynkaruk, A.P. Prout, K. Nygard, R. Veldhuizen, R. Hammond, B.S Richardson

## Abstract

Supported by grants from the Canadian Institute of Health Research (CIHR) and Lawson Health Research Institute (LHRI) Internal Research Fund (MGF and BSR); CIHR, Fonds de la recherche en santé du Québec (FRSQ) (MGF). BSR is the recipient of the Canada Research Chair in Fetal and Neonatal Health and Development.

Neuroinflammation *in utero* may contribute to brain injury resulting in life long neurological disabilities. The pivotal role of the efferent cholinergic anti-inflammatory pathway (CAP) in controlling inflammation has been described in adults, but its importance in the fetus is unknown. Moreover, it is unknown whether CAP may also exert anti-inflammatory effects on the brain via CAP’s afferent component of the vagus nerve. Based on multiple clinical studies in adults and our own work in fetal autonomic nervous system, we gauged the degree of CAP activity *in vivo* using heart rate variability measures reflecting fluctuations in vagus nerve activity. Measuring microglial activation in the ovine fetal brain near-term, we show *in vivo* that afferent fetal CAP may translate increased vagal cholinergic signaling into suppression of cerebral inflammation in response to near-term hypoxic acidemia as might occur during labour. Our findings suggest a new control mechanism of fetal neuroinflammation via the vagus nerve, providing novel possibilities for its non-invasive monitoring *in utero* and for targeted treatment.

## INTRODUCTION

Induced animal sepsis and clinical-pathologic studies in adults indicate that loss of the cholinergic anti-inflammatory pathway’s (CAP) inhibitory influence unleashes innate immunity, producing higher levels of pro-inflammatory mediators that exacerbate tissue damage. This decrease in CAP activity also decreases short-term heart rate variability (HRV), *e.g.*, as measured by the beat-to-beat HRV measures, such as root mean square of successive differences in R-R intervals of ECG (RMSSD), a measure of vagal modulation of HRV. [1, 2] Thus, short-term HRV measures reflect CAP activity in adults. [3] Of note, RMSSD also reflects vagal activity in fetal sheep. [4]

Increased CAP vagal activity inhibits the release of pro-inflammatory cytokines such as interleukin (IL)-1β. [1] This systemic CAP effect is mediated via the α7 nicotinic acetylcholine receptor (α7nAChR) expressed on macrophages. [5] However, recent studies have shown a similar α7nAChR-dependent effect in brain microglia *in vitro*. [6, 7,8]

In adult species, high-mobility group box protein 1 (HMGB1), a non-histone DNA-binding protein, acts as an important pro-inflammatory cytokine linking necrosis with ensuing inflammation by translocating from the neuronal nucleus to the cytosol and then to the extracellular space, leading to microglial activation. [9] Much attention has been paid to the effects of α7nAChR stimulation on HMGB1 secretion because of its therapeutic potential to treat sepsis; HMGB1 represents a crucial link between neuronal necrosis and the cerebral inflammatory response mediated by microglia, thus impacting the long term outcome of neurological injury. [10, 9] HMGB1 also acts as a potent pro-inflammatory cytokine when secreted by microglia in response to inflammatory stimuli. [11] This requires translocation of HMGB1 from nucleus to cytosol. [9]

Systemic and neuroinflammation have been implicated as important pathophysiological mechanisms acting independently to cause fetal brain injury or contributing to hypoxic-asphyxial brain injury with consequences for postnatal health. [12,13] In the late-gestation ovine and human fetus, the autonomic nervous system and cholinergic vagal activity in particular are known to be sufficiently mature. [2,14]

We have shown that CAP is active spontaneously near term, such that individual baseline RMSSD values and the levels of the pro-inflammatory cytokines IL-1β and IL-6 are inversely correlated, reflecting spontaneous CAP activity. [15]

First, we hypothesized that the fetal inflammatory response induced by hypoxic-acidemia will result in an increase of systemic CAP activity as a compensatory mechanism and an inhibitory effect of CAP on the cerebral inflammatory response. The systemic inflammatory response will be reflected by an increased vagal activity and hence a correlation of RMSSD and IL-1β.

Second, we sought to determine the effect of fetal hypoxic-acidemia insult on brain regional activation of the microglia expressing α7nAChR, and the relation of systemic and cerebral CAP activation to the intracellular HMGB1 localization in these cells. Thus, we hypothesized that the cerebral inflammatory response will result in microglial HMGB1 translocation from the nucleus to the cytosol due to increased microglial activation via α7nAChR and this HMGB1 translocation will correlate with the degree of CAP’s vagal activation measured by RMSSD.

## RESULTS

As reported, repetitive UCO resulted in worsening acidosis over 3 to 4 h and eventually a severe degree of acidemia, fetal pH 7.36 ± 0.03 to 6.90 ± 0.13 (p<0.01). [16]

RMSSD and IL-1β increased ~2 fold from baseline versus the time of nadir pH (p < 0.05), and fell by 1 h of recovery (Fig. 2A, 2B; for IL-1β cf. [17]). Of note, at 1 h of recovery the values of RMSSD and IL-1β were still clearly, but not statistically significantly elevated. Baseline, nadir pH and 1 h of recovery RMSSD correlated to corresponding IL-1β levels at R = 0.57 (p=0.02, n=17, Fig. 2C).

Fetal gender did not contribute to this correlation. As reported, MG cell counts were increased in the white matter of the treated animals versus the control group. [17] RMSSD measurements at baseline and 1 h of recovery correlated inversely to white matter MG cell counts determined at 24 h of recovery at R = −0.71 (p=0.05) and R = −0.89 (p=0.03), respectively (Fig. 2D).

**Figure 1.**
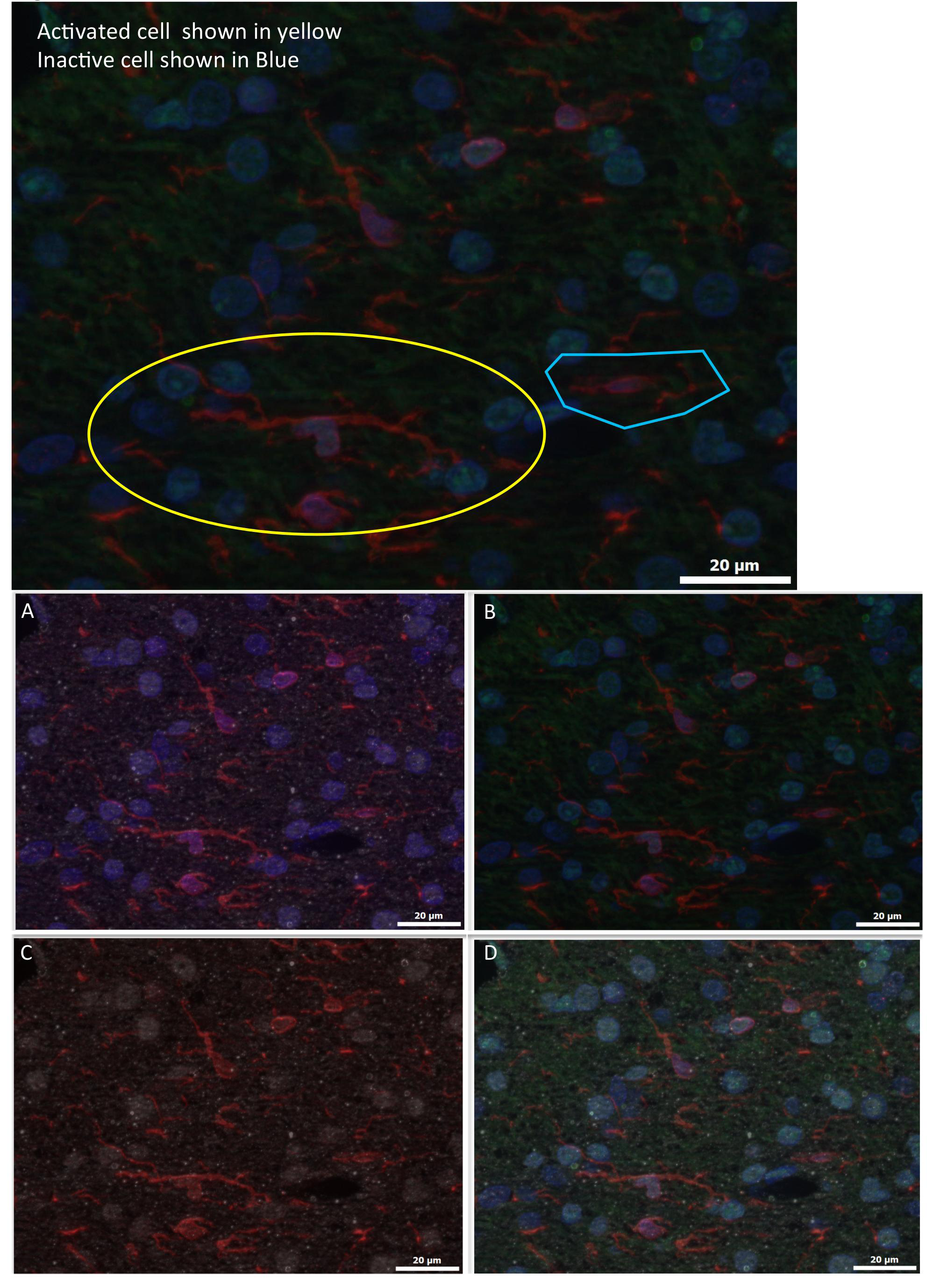
α7nAChR+HMGB1+Iba1+DAPI immunofluorescent staining: Red=Iba1; Blue=DAPI;Green=HMGB1; White=α7nAChR. Images were made from extended depth of focus, maximum intensity projections from convexial grey matter layers 4 to 6. **TOP:** Features of active versus inactive microglia: Inactive cells have no interaction with neurons, are smaller, less bright and less ramified versus the activated cells. **BOTTOM:** Co-localization staining for Iba1+ cells expressingα7nAChR with intracellular localization of HMGB1 and DAPI counterstain. **A**: Iba1+DAPI+α7nAChR; **B**: Iba1+DAPI+HMGB1; **C**: Iba1+ α7nAChR; **D**: Best focus color composite of Iba1+ α7nAChR+HMGB1+DAPI.

**Figure 2.**
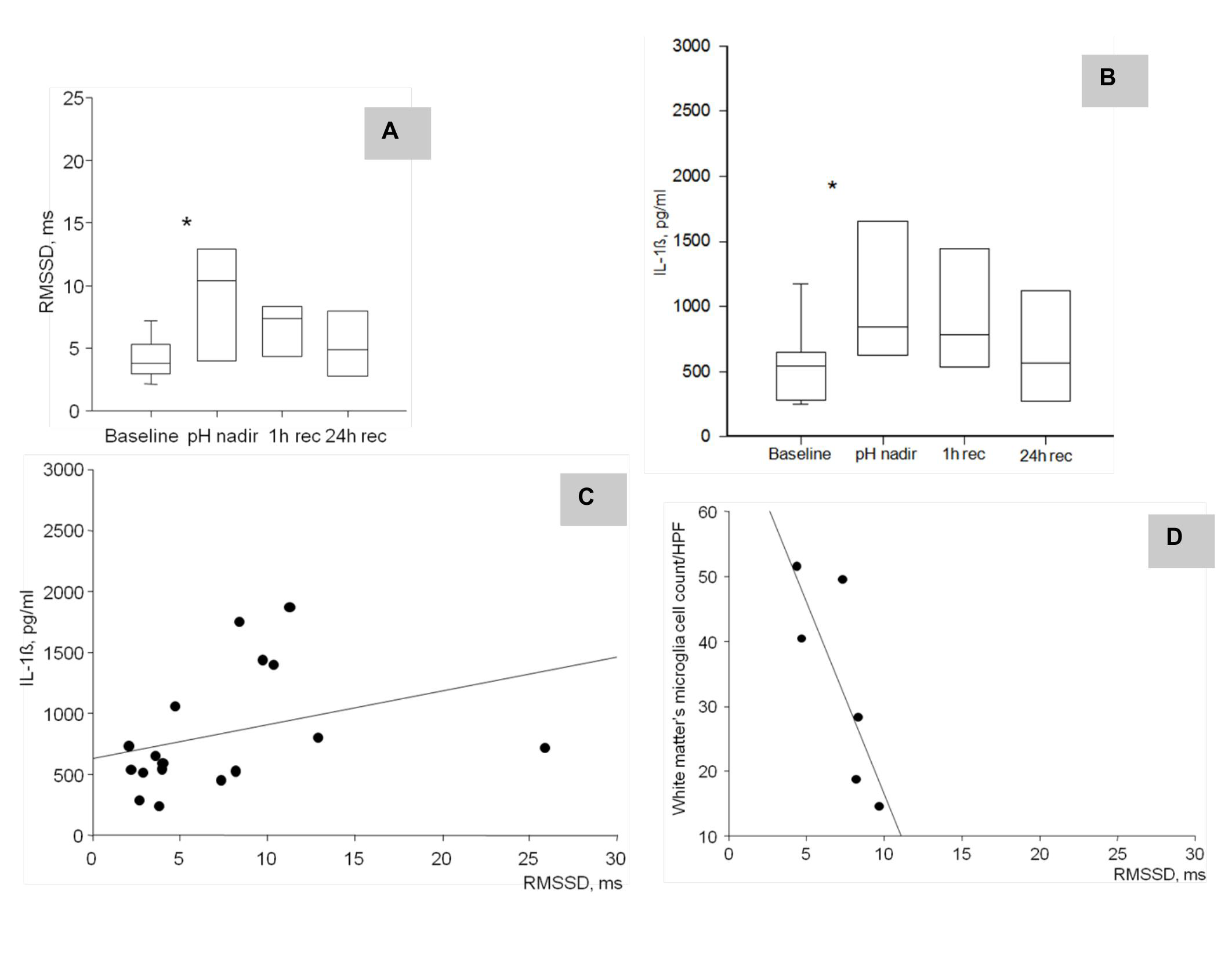
**A and B.** Fetal inflammatory response to acute hypoxic-acidemia. * p<0.05 versus baseline.pH nadir, at pH<7.00; recovery 1 and 24 hours after the pH nadir. Median±{5-95%}. **C.** IL-1β measured at baseline, pH nadir and 1 h or recovery correlates to RMSSD (R=0.57, p=0.02, n=17). Baseline values were chosen for lactate >1.5 mmol/l. **D.** White matter microglia cell counts at 24 h recovery correlate to RMSSD at 1h of recovery (R=0.89, p=0.03, n=6).

For both HMGB1 translocation index and α7nAChR expression, the main effect of belonging to the UCO or the control group was not significant (p=0.25 and p=0.54, respectively, Fig. 3). However, the effect of group on HMGB1 translocation differed according to brain region (mostly in hippocampus and GM46) and microglia status (p<0.001 for interaction terms) (Table 1). Interestingly, statistically significant interaction effects on HMGB1 translocation were observed in cortical and hippocampal regions, but not in the subcortical (thalamus, white matter) brain regions. In grey matter, these effects applied to quiescent MG (qMG), but not to activated MG (aMG); meanwhile, in hippocampus this was mostly apparent in aMG.

**Table 1.** Effect of UCO, microglia status and brain regions on the HMGB1 translocation index. Parameter Estimates

| Parameter | B | Std. Error | 95% Wald Confidence Interval |  | Hypothesis Test |  |  |
| --- | --- | --- | --- | --- | --- | --- | --- |
|  |  |  | Lower | Upper | Wald Chi-Square | df | Sig. |
| (Intercept) | .535 | .0646 | .408 | .661 | 68.416 | 1 | .000 |
| [Group=UCO ] | .167 | .0793 | .012 | .322 | 4.445 | 1 | .035 |
| [Group=Control ] | 0<sup>a</sup> | . | . | . | . | . | . |
| [Group=UCO ] * [Brain_region=CA1 ] * [Microglia_status=active ] | .073 | .0675 | -.059 | .205 | 1.173 | 1 | .279 |
| [Group=UCO ] * [Brain_region=CA1 ] * [Microglia_status=quiescent ] | .090 | .0907 | -.088 | .268 | .989 | 1 | .320 |
| **[Group=UCO ] \* [Brain_region=CA3 ] \* [Microglia_status=active ]** | .106 | .0536 | .001 | .211 | 3.931 | 1 | .047 |
| [Group=UCO ] * [Brain_region=CA3 ] * [Microglia_status=quiescent ] | .019 | .0490 | -.077 | .115 | .149 | 1 | .699 |
| **[Group=UCO ] \* [Brain_region=DG ] \* [Microglia_status=active ]** | .182 | .0445 | .095 | .270 | 16.795 | 1 | .000 |
| [Group=UCO ] * [Brain_region=DG ] * [Microglia_status=quiescent ] | .102 | .0809 | -.057 | .260 | 1.589 | 1 | .207 |
| [Group=UCO ] * [Brain_region=GM13 ] * [Microglia_status=active ] | -.102 | .0727 | -.244 | .041 | 1.951 | 1 | .163 |
| **[Group=UCO ] \* [Brain_region=GM13 ] \* [Microglia_status=quiescent ]** | -.203 | .0712 | -.342 | -.063 | 8.112 | 1 | .004 |
| [Group=UCO ] * [Brain_region=GM46 ] * [Microglia_status=active ] | -.062 | .0648 | -.189 | .065 | .902 | 1 | .342 |
| **[Group=UCO ] \* [Brain_region=GM46 ] \* [Microglia_status=quiescent ]** | -.125 | .0593 | -.241 | -.009 | 4.451 | 1 | .035 |
| [Group=UCO ] * [Brain_region=Thalamus] * [Microglia_status=active ] | -.051 | .0459 | -.141 | .039 | 1.219 | 1 | .270 |
| [Group=UCO ] * [Brain_region=Thalamus] * [Microglia_status=quiescent ] | -.052 | .0553 | -.160 | .056 | .889 | 1 | .346 |
| [Group=UCO ] * [Brain_region=WM ] * [Microglia_status=active ] | -.030 | .0234 | -.076 | .016 | 1.592 | 1 | .207 |
| [Group=UCO ] * [Brain_region=WM ] * [Microglia_status=quiescent ] | 0<sup>a</sup> | . | . | . | . | . | . |
| **[Group=Control ] \* [Brain_region=CA1 ] \* [Microglia_status=active ]** | .117 | .0490 | .021 | .213 | 5.718 | 1 | .017 |
| **[Group=Control ] \* [Brain_region=CA1 ] \* [Microglia_status=quiescent ]** | .254 | .0777 | .102 | .406 | 10.677 | 1 | .001 |
| **[Group=Control ] \* [Brain_region=CA3 ] \* [Microglia_status=active ]** | .177 | .0811 | .018 | .336 | 4.755 | 1 | .029 |
| **[Group=Control ] \* [Brain_region=CA3 ] \* [Microglia_status=quiescent ]** | .130 | .0584 | .015 | .244 | 4.933 | 1 | .026 |
| **[Group=Control ] \* [Brain_region=DG ] \* [Microglia_status=active ]** | .118 | .0548 | .010 | .225 | 4.621 | 1 | .032 |
| [Group=Control ] * [Brain_region=DG ] * [Microglia_status=quiescent ] | .057 | .0563 | -.053 | .168 | 1.037 | 1 | .309 |
| [Group=Control ] * [Brain_region=GM13 ] * [Microglia_status=active ] | .088 | .0937 | -.095 | .272 | .891 | 1 | .345 |
| [Group=Control ] * [Brain_region=GM13 ] * [Microglia_status=quiescent ] | .049 | .1170 | -.180 | .278 | .175 | 1 | .676 |
| [Group=Control ] * [Brain_region=GM46 ] * [Microglia_status=active ] | .142 | .0932 | -.041 | .325 | 2.318 | 1 | .128 |
| **[Group=Control ] \* [Brain_region=GM46 ] \* [Microglia_status=quiescent ]** | .228 | .0913 | .049 | .407 | 6.253 | 1 | .012 |
| [Group=Control ] * [Brain_region=Thalamus] * [Microglia_status=active ] | .106 | .0704 | -.032 | .244 | 2.250 | 1 | .134 |
| [Group=Control ] * [Brain_region=Thalamus] * [Microglia_status=quiescent ] | .112 | .1129 | -.110 | .333 | .975 | 1 | .323 |
| [Group=Control ] * [Brain_region=WM ] * [Microglia_status=active ] | .001 | .0354 | -.069 | .070 | .001 | 1 | .981 |
| [Group=Control ] * [Brain_region=WM ] * [Microglia_status=quiescent ] | 0<sup>a</sup> | . | . | . | . | . | . |
| (Scale) | .025 | . | . | . | . | . | . |

Dependent Variable: HMGB1_translocation Model: (Intercept), Group, Group * Brain_region * Microglia_status (*active* or quiescent) a. Set to zero because this parameter is redundant. GM13 and 46 are cortical grey matter layers 1-3 and 4-6, respectively; WM, white matter; CA1, CA3 and DG (dentate gyrus) are the hippocampal subregions **Bold** entries are statistically significant results.

**Figure 3.**
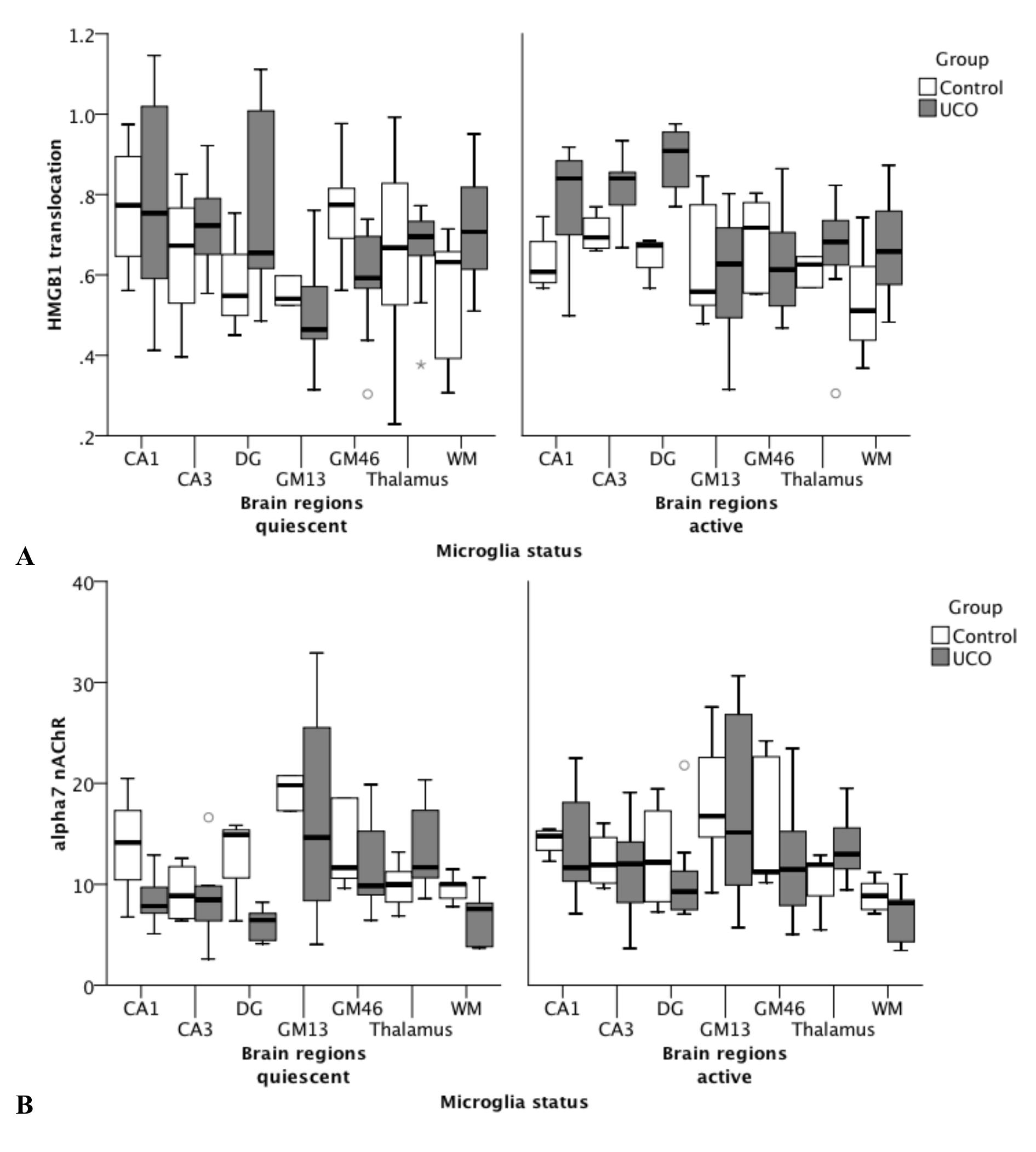
**A.** Effect of UCO, microglia status and brain regions on HMGB1 translocation: significantgroup*brain region*microglia status interaction (p<0.001). See Table 1 for details. **B.** Effect of UCO and microglia status on α7 nAChR immunofluorescence measured as intensity per area: significant group*microglia status*HMGB1 translocation interaction (p<0.001). See Table 2 for details. Note that, for α7 nAChR signal, between-brain regions comparisons were not possible, because gain settings were optimized for each brain region and kept constant between cell compartment and groups (but not from region to region). HMGB1 signal is expressed as ratio of cytosolic to nuclear signal, *i.e.*, the higher the ratio, the more HMGB1 translocation is observed; this normalization permits between-brain regions comparisons.

Similarly, a model that accounted for interactions of group and microglia status and HMGB1 was able to predict α7nAChR expression (p<0.001). Notably, the effect of UCO group was in the same direction but of much greater magnitude in aMG compared to qMG (Table 2).

**Table 2.** Effect of UCO, microglia status and HMGB1 translocation on α7 nAChR signal Parameter Estimates

| Parameter | B | Std. Error | 95% Wald Confidence Interval |  | Hypothesis Test |  |  |
| --- | --- | --- | --- | --- | --- | --- | --- |
|  |  |  | Lower | Upper | Wald Chi-Square | df | Sig. |
| (Intercept) | 15.224 | 5.1517 | 5.127 | 25.321 | 8.733 | 1 | .003 |
| [Group=UCO ] | 3.842 | 6.2720 | -8.451 | 16.135 | .375 | 1 | .540 |
| [Group=Control ] | 0<sup>a</sup> | . | . | . | . | . | . |
| **[Group=UCO ] \* [Microglia_status=active ] \* HMGB1_trans** | -9.527 | 4.5915 | -18.527 | -.528 | 4.306 | 1 | .038 |
| **[Group=UCO ] \* [Microglia_status=quiescent ] \* HMGB1_trans** | -12.360 | 4.5838 | -21.344 | -3.376 | 7.271 | 1 | .007 |
| [Group=Control ] * [Microglia_status=active ] * HMGB1_trans | -3.069 | 8.2511 | -19.241 | 13.103 | .138 | 1 | .710 |
| [Group=Control ] * [Microglia_status=quiescent ] * HMGB1_trans | -3.264 | 7.9857 | -18.915 | 12.388 | .167 | 1 | .683 |
| (Scale) | 35.371 |  |  |  |  |  |  |

Dependent Variable: alpha7nAChR Model: (Intercept), Group, Group * Microglia_status (*active* or quiescent) * HMGB1_translocation a. Set to zero because this parameter is redundant. **Bold** entries are statistically significant results.

In parallel, RMSSD at 1 hour of recovery correlated highly with cytosolic HMGB intensity per area in aMG of thalamus (R = −0.94, p=0.005, Fig. 4A) and RMSSD at pH nadir correlated with α7nAChR intensity per area in aMG of WM (R = 0.83, p=0.04, Fig. 4B). Similar to the relationship shown in Fig 4A, within both qMG and aMG in WM, HMGB1 translocation index correlated to RMSSD at 1 hour of recovery (R = −0.83, p=0.04 and R = −0.89, p=0.02, respectively). This finding was again replicated for qMG of GM13 and aMG of GM46: HMGB1 translocation index correlated there to RMSSD at pH nadir (R = −0.99, p<0.001 and R = −0.83, p=0.04, respectively). That is, higher RMSSD values correlated with lower HMGB1 translocation and higher α7nAChR intensity per area in brain region-specific and microglia status-specific manner.

**Figure 4.**
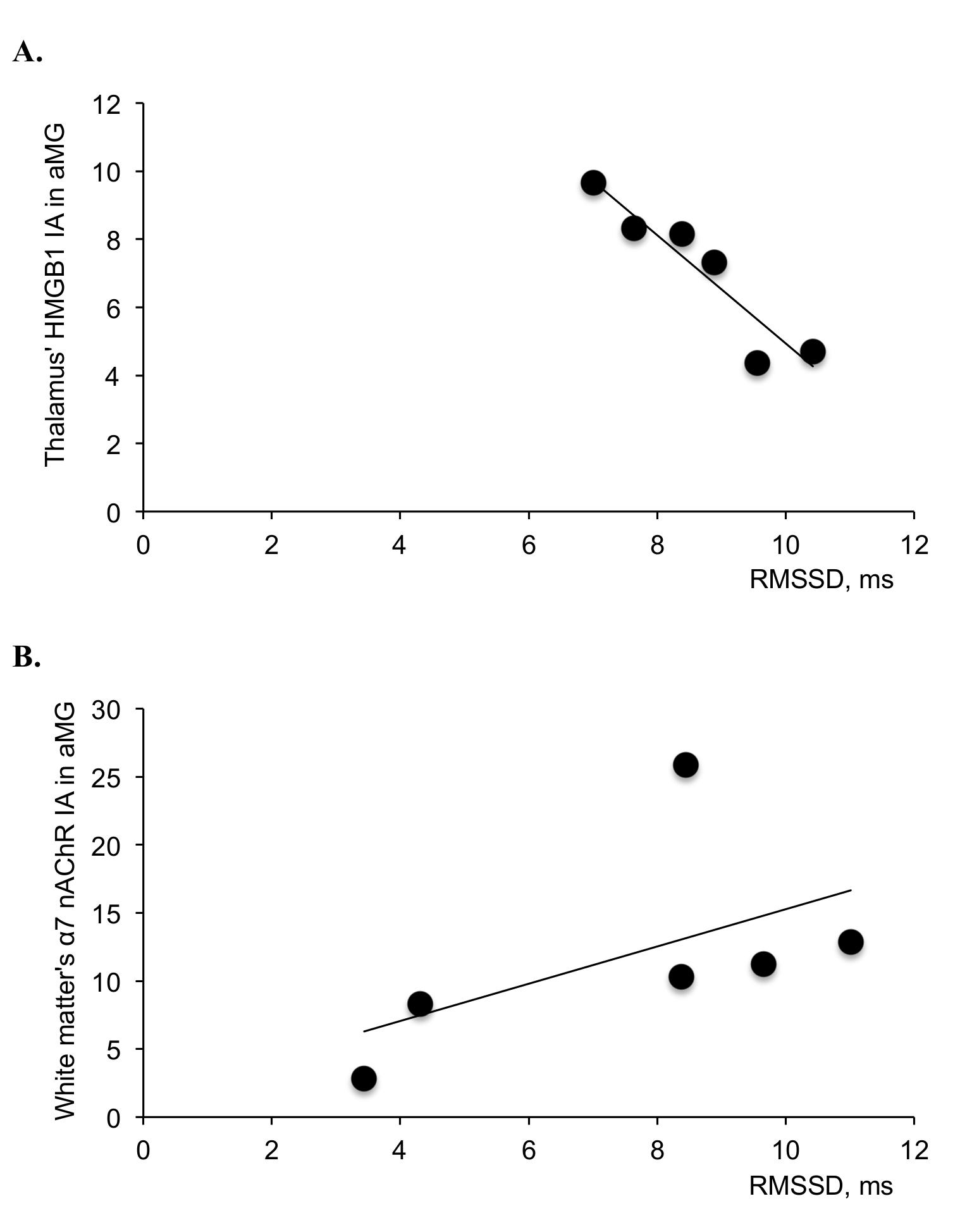
**UCO group:** HMGB1 and α nAChR immunofluorescence are shown, measured as intensities per area (I/A) in activated microglia (aMG) in relation to RMSSD as marker of CAP activity. RMSSD at 1 hour recovery negatively correlates with cytoplasmic HMGB1 in thalamus (**A**), R= −0.94, p=0.005. RMSSD at pH nadir positively correlates with α7 nAChR in the white matter (**B**), R=0.83, p=0.04.

## Discussion

We demonstrate that UCO-induced insult simulating human labour results in increasing fHRV properties known to reflect fluctuations of vagal activity which are correlated with systemic and brain inflammatory responses and with shifts in nucleus-cytosol HMGB1 distribution in α7AChRs-positive microglia. We propose that this behaviour is due to an increase in afferent CAP activity exerting anti-neuroinflammatory effects via microglial α7AChRs by limiting microglial activation. We observe that such neuroprotective effects occur in a brain region-dependent manner.

### Efferent effects of fetal CAP and its relation to HRV

Our findings support the first hypothesis that near-term asphyxia with worsening acidemia leads to vagal activation correlated to the degree of systemic inflammatory response, which suggests CAP activation. [4,16,17] Fetal acidemia, but not intermittent chronic hypoxia alone, seems to be required for an aseptic induction of fetal inflammatory response and activation of CAP. [17] Consequently, the inhibition of brain regional cellular innate immune response was found in fetuses that were subject to acute worsening acidemia, but not to chronic intermittent hypoxia. Of note, duration of UCO insults, but not levels of lactate per se correlated to IL-1β in the near-term fetuses. [17] In parallel with positive correlation of IL-1β and RMSSD, this suggests that chemoreceptor-mediated vagal activation results in part from the CAP-mediated sensing of the systemic inflammatory response that is more pronounced with longer duration of the stimulus.

### Afferent effects of fetal CAP on neuroinflammation via α7nAChR carrying microglia

To our knowledge this is the first report relating systemic vagal activity *in vivo* and the cerebral inflammatory response in α7nAChR-labeled microglia, thus providing *in situ* and *in vivo* support for our second hypothesis, namely that CAP not only acts via efferent peripheral pathway to suppress systemic levels of inflammation, but also targets neuroinflammation in the brain. This anti-inflammatory effect is probably achieved via the afferent branch of vagus nerve carrying cholinergic signalling to the brainstem where it is relayed to the higher subcortical and cortical regions. This notion is supported by several considerations. First, ~80% of vagus nerve fibers are afferent and may provide a retrograde feedback to the cerebral CAP system synchronizing its responses with those of the systemic CAP activity to the endogenous (*e.g.*, acidemia as shown in the present study) or exogenous (*e.g.*, LPS, bacteria or viruses) inflammatory stimuli. [18] Second, vagus nerve stimulation (VNS) has been used to treat drug-refractory epilepsy for decades, yet the mechanism of action for VNS has remained elusive. [19] Anti-inflammatory effects of VNS may play a part in resetting the brain’s epileptogenic potential. [19] Third, recent nascent work on mapping the neuroimmunological homunculus has provided further evidence for the far-reaching connection between peripheral inflammation and cerebral perception and responses that are likely to include the vagus nerve as one of the main communication carriers both to and from the brain. [20, 21] Fourth, unlike its other cholinergic counterparts, the α7nAChR can be activated by a primary ligand other than acetylcholine, *i.e.,* choline. [22] Hence the α7nAChR can function in areas of the brain devoid of cholinergic transmission *per se*, where the far more ubiquitous choline may act as a substitute ligand. This may further contribute to a widespread effect of cholinergic signalling in the brain via the α7nAChR.

Furthermore, we demonstrate *in situ* α7nAChR expression in fetal sheep microglia and the modulation of this receptor’s immunofluorescent signal from nucleus and cytosol/membrane compartments by hypoxic-acidemia and microglia status. We show that fetuses with higher RMSSD during severe acidemia showed lower MG activation in the WM. We further show that this may be mediated via α7nAChR in the WM's activated MG. α7nAChR stimulation on adult and neonatal murine derived microglia *in vitro* reduces secretion of LPS-induced pro-inflammatory cytokine TNF-α. [23, 24] The CAP-mediated effect of afferent cholinergic activity on microglial HMGB1 release is further supported by negative correlation between the microglial thalamic cytosolic HMGB1 and vagal activation after reaching most severe acidosis. The α7nAChR expression is not static within a given brain region, but rather appears to be dynamically regulated by stimuli such as hypoxic-ischemic injury which down-regulates α7nAChR mRNA, unless the receptor is agonistically stimulated. [7] We propose that such stimulation may not only be exogenous, pharmacologically driven, but also endogenous and occurring via afferent CAP. Indeed, fetal cerebral inflammatory response and ensuing CAP activation resulted in a pronounced microglial α7nAChR intensity per area increase in the periventricular WM and reduced cytoplasmic HMGB1 intensity per area in the thalamus, which suggests that CAP homeostatically modulates microglia activity towards a neuroprotective phenotype via α7nAChR, similar teleologically to what has been proposed by Tracey *et al.* for systemic peripheral efferent effects of CAP. [5]

We identified microglia status-dependent differences in HMGB1 translocation between cortical and subcortical brain regions. These differences seemed amplified by the UCO insult. UCOs are known to induce a redistribution of the regional cerebral blood flow (rCBF) toward subcortical structures. [25] In the hippocampal subregions, HMGB1 translocation was noted in the active microglia, while in the cortical layers 1-6 this was true for the quiescent microglia. This may be explained by a reportedly stronger rCBF increase in hippocampus than in cortex under a similar UCO insult. [25] The relative rCBF increase in hippocampus may also increase its exposure to the systemically circulating pro-inflammatory cytokines and lactic acid which up regulates microglia, in our approach captured by a stronger Iba1+ signal, larger microglial cell bodies and processes with close proximity to the neurons. [26, 27] This finding further contributes to the observation that hippocampus is one of the brain regions most susceptible to hypoxic-ischemic insults. [25] Further studies are needed to better characterize the microglia phenotypes involved in the UCO insult.

### Clinical implications

In a series of experiments in chronically instrumented unanaesthetized fetal sheep, an important model of human pregnancy, we showed that a fetal systemic and cerebral inflammatory response required the presence of severe hypoxic-acidemia and was not induced by chronic hypoxia alone as might occur inthe late gestation human fetus during labour or antenatally with growth restriction, respectively. [28] Since RMSSD, as a measure of vagal activity, increases in response to perinatal acidemia and inflammation, [4,29] we sought to determine the extent to which measures of systemic and cerebral inflammation in the fetus relate to RMSSD and indicate peripheral and cerebral CAP activity. We have induced fetal inflammation with hypoxic-acidemia. [30] This stimulus activates systemic and cerebral innate immune responses and corresponds to a frequently observed spectrum of fetal heart rate distress patterns during human labour. Severe fetal acidemia (pH<7.00) at birth is observed in ~0.5-10% of human births. [31, 32] Approximately 20% of these babies will have neurologic sequelae including hypoxic-ischemic encephalopathy (HIE) and cerebral palsy. [33-35] Every 5^th^ baby with cerebral palsy will have had an asphyxial event during birth, and every 10^th^, some degree of inflammatory exposure, with antenatal growth restriction and chronic hypoxia identified as another major contributing factor.

This is supported by animal research indicating that chronic hypoxia, prior to an acute asphyxia event, alters the interplay between the fetal neuroinflammatory and neuronal responses to the insult, potentially exacerbating the long-term effects of the exposure. [26] Overall, antenatal hypoxia and perinatal hypoxic-acidemia are major contributors to perinatal brain injury resulting in increased risk for acute or life-long morbidity and mortality. [37, 38] Our findings suggest that an endogenous neuroinflammatory control mechanism, CAP, plays a neuroprotective role in etiology of early perinatal brain injury.

### Conclusions

Our study provides first evidence of a neuroimmunological link in the late gestation fetus. Enhancing fetal CAP activity may suppress activation of microglia, therefore promoting neuroprotection. This may improve postnatal short- and long-term health outcomes through decreasing lasting brain injury.

## Methods

### Ethics Statement

This study was carried out in strict accordance with the recommendations in the Guide for the Care and Use of Laboratory Animals of the National Institutes of Health. The protocol has been approved by the Committee on the Ethics of Animal Experiments of the University of Western Ontario (Permit Number 2006-091-08 “Intrapartum fetal monitoring: markers of hypoxic related injury”).

### Surgical Preparation

Fetal sheep near term (0.86 gestation) of mixed breed were surgically instrumented with umbilical cord occluders to induce hypoxic-acidemia (n=10+5, further referred to as *UCO group* compared to *Control group* which was also instrumented but not subjected to UCOs). The anesthetic and surgical procedures and postoperative care of the animals have been described (23). Briefly, using sterile technique under general anesthesia (1g thiopental sodium in solution intravenously (IV) for induction; Abbott Laboratories Ltd., Montreal, Canada; followed by 1% to 1.5% halothane in O_2_ for maintenance), a midline incision was made in the lower abdominal wall, and the uterus was palpated to determine fetal number and position. The upper body of the fetus and proximal portion of the umbilical cord were exteriorized through an incision in the uterine wall. Polyvinyl catheters (Bolab, Lake Havasu City, AZ) were placed in the right and left brachiocephalic arteries, and the right brachiocephalic vein. Stainless steel electrodes were implanted biparietally on the dura for the recording of electrocortical activity (ECoG) and over the sternum for recording electrocardiogram (ECG). An inflatable silicone occluder cuff (OCHD16; In Vivo Metric, Healdsburg, CA) was positioned around the proximal portion of the umbilical cord and secured to the abdominal skin. Once the fetus was returned to the uterus, a catheter was placed in the amniotic fluid cavity and subsequently in the maternal femoral vein. Antibiotics were administered intra-operatively to the mother, (0.2 g trimethoprim and 1.2 g sulfadoxine, Schering Canada Inc., Pointe-Claire, Canada), fetus and amniotic cavity (1 million IU penicillin G sodium, Pharmaceutical Partners of Canada, Richmond Hill, Canada). Amniotic fluid lost during surgery was replaced with warm saline. The uterus and abdominal wall incisions were sutured in layers and catheters exteriorized through the maternal flank and secured to the back of the ewe in a plastic pouch.

Animals were allowed a 3-4 day postoperative period prior to experimentation, during which the antibiotic administration was continued. Arterial blood was sampled each day for evaluation of fetal condition and catheters were flushed with heparinized saline to maintain patency.

### Data and blood sample acquisition

A computerized data acquisition system was used to record pressures in the fetal brachiocephalic artery and amniotic cavity at 256 Hz, and the ECG and ECoG signals at 1000 Hz. All signals were monitored continuously throughout the experiment using Chart 5 for Windows (ADInstruments Pty Ltd, Bella Vista, Australia).

Fetal 3 mL blood samples were immediately spun at 4°C (4 minutes, 4000 rpm; Beckman TJ-6, Fullerton, CA) and the plasma decanted and stored at −80°C for subsequent cytokine analysis. In addition, to characterize baseline health status of the animals fetal 1 mL blood samples were analyzed for blood gas values, pH, glucose and lactate with an ABL-725 blood gas analyzer (Radiometer, Copenhagen, Denmark) with temperature corrected to 39°C.

At the end point of each experiment as detailed below, the ewe and fetus were killed with an overdose of barbiturate (30 mg pentobarbital sodium, Fatal-Plus; Vortech Pharmaceuticals, Dearborn, MI) and a post mortem was carried out during which fetal gender and weight were determined, and the location and function of the umbilical cord occluder cuff were confirmed. The fetal brain was then perfusion-fixed with 500 mL of cold saline followed by 500 mL of 4% paraformaldehyde and processed for histochemical analysis as reported. [39], [16, 40]

### Experimental procedures

As reported [17, 41], animals were studied through a 1 to 2 hour baseline period, an experimental period of repetitive UCO with worsening acidemia, and were then allowed to recover overnight. Five additional age-matched additional animals were used as controls for immunohistochemistry.

After the baseline period which began at ~8 a.m., repetitive UCO were performed with increasing frequency until severe fetal acidemia was detected (arterial pH<7.00), at which time the UCO were stopped. Complete UCO was induced by inflation of the occluder cuff with ~5 mL saline solution, the exact volumes having been determined by visual inspection and testing at the time of surgery for each animal. During the first hour a mild UCO series was performed consisting of cord occlusion lasting for 1 minute and repeating every 5 minutes. During the second hour a moderate UCO series was performed consisting of cord occlusion for 1-minute duration and repeating every 3 minutes. During the third hour a severe UCO series was performed consisting of cord occlusion for 1 minute duration, repeated every 2 minutes, and this series was continued until the targeted fetal arterial pH was attained. Following the mild as well as the moderate UCO series 10 minute periods with no UCO were undertaken, during which fetal arterial blood was sampled and arterial blood pressure, ECOG, and ECG data were recorded in the absence of fetal heart rate decelerations. After attaining the targeted fetal arterial pH of <7.00 and stopping the repetitive UCO, animals were allowed to recover for ~24 hours.

Fetal arterial blood samples were obtained during the baseline period (3 mL), at the end of the 1^st^ UCO of each UCO series (1 mL), and ~5 minutes after each UCO series (3 mL). In addition, fetal arterial blood samples were obtained between UCO at ~20 and 40 minutes of the moderate and severe UCO series (1 mL), and at 1, 2 and 24 hours of recovery (3 mL). At ~4 p.m. on day 2, the animals were killed as described above.

### Measurements of inflammatory responses

#### Systemic IL-1β and IL-6 secretion

An ELISA was used to analyse in duplicate the concentrations of IL-1β and IL-6 in fetal arterial and maternal venous plasma samples as reported. [17] IL-1β and IL-6 standards were purchased from the University of Melbourne, Centre of Animal Biotechnology, Melbourne, Australia. Mouse anti-ovine IL-1β (MAB 1001) and IL-6 (MAB 1004) antibodies and rabbit anti-ovine IL-1β (AB 1838) and IL-6 (AB 1889) polyclonal antibodies were purchased from Chemicon International, Temecula, CA. Separate 96-well plates were coated with mouse monoclonal ovine IL-1β or IL-6 antibody (1:200, in 0.1M NaCO_3_, pH to 9.6) and incubated overnight at 4°C. The following day, plates were washed three times with wash solution (1X PBS with 0.05%Tween, pH to 7.4) to remove excess monoclonal antibody. Plates were then blocked with assay diluent (555213, BD OptEIA, BD Biosciences) at room temperature for 1 hour. Wells were then rinsed three times with the wash solution followed by aliquoting standards (40,000 pg/mL to 156 pg/mL and blanks) and samples, and incubation on the shaker at room temperature for 2 hours. Subsequently, wells were rinsed three times with washing solution and the appropriate rabbit anti-ovine polyclonal antibody (IL-1β or IL-6, 1:500) was added to each well and incubated on the shaker for 1 hour. Following five more washes, HRP-donkey anti rabbit IgG (AP182p, Chemicon International, 1:10,000) was added to each well and incubated on the shaker for 1 hour. The wells were then washed seven times with wash solution to remove all unbound secondary antibody, followed by the addition and 30 minute incubation with substrate solution (51-2606KC & 51-2607KC, BD Biosciences) in the dark. Stop solution (1N H_2_SO_4_) was applied and each well was read using a spectrophotometer at 450 nm, with 575 nm wavelength correction.

### Immunohistochemistry: Microglia counts

These methods and findings have been reported. [17] The focus of this study was the putative relationship between the changes in microglia (MG) count in the white matter we had reported and the fHRV as proxy for CAP activity. Below we report the method as it was applied for all brain regions. Thepresence of MG in brain tissue sections was determined by avidin-biotin-peroxidase complex enhanced immunohistochemistry (Vectastain Elite; Vector Laboratories, Burlingame, CA) as previously reported. [17, 40] To reduce staining variability, all immunohistochemistry was performed on the same day with the same batch of antibody and solutions. Tissue sections were incubated with an anti-IBA1 rabbit polyclonal antibody (1:500, Wako Industries, Richmond, VA), a robust marker for sheep MG, with detection of bound antibody obtained following incubation in Cardassian DAB Chromogen (Biocare Medical, Concord, CA)

Brain regions that were selected from each animal for analysis were taken from a coronal section of blocked cerebral hemisphere tissue at the level of the mammillary bodies and included the parasagittal and convexity cerebral gray matter, periventricular white matter, thalamus, CA1, dentate gyrus (DG) and the combined CA2 and CA3 regions of the hippocampus. Each of the parasagittal and convexity cerebral gray matter regions was further divided into sub-regions combining layers 1, 2, and 3 and layers 4, 5, and 6. Image analysis was performed with a transmitted light microscope (Leica DMRB, Leica-Microsystems, Wetzler, Germany) at 40x magnification. Positive MG cell immunostaining was quantified with an image analysis program (Image Pro Plus 6.0, Media Cybernetics, Silver Spring, MD). The image analysis system was first calibrated for the magnification settings that were used, and thresholds were established to provide even lighting and no background signal. Six high-power field (HPF) photomicrographs (HPF area = 7cm^2^) per brain region/subregion per animal were randomly collected as a 24 bit RGB colour modeled image. The same illumination setting was applied to all images for all of the brain regions. Using the Image Pro Plus’ RGB colour range selection tool, colour samplings of positive DAB stained areas were obtained from multiple brain regions and tested for specificity against the negative control. Appropriate ranges of colour were selected showing positive contiguous cytoplasmic staining as a criterion for MG cell count scoring, which were then applied uniformly to calibrated images for all brain regions. Scoring was then performed automatically in a blinded fashion to experimental groups.

### Immunohistochemistry: HMGB1 translocation in Iba1+ microglia expressing α7 nAChR

Similar to MG analyses, whole brain coronal sections through the level of the mammillary body were selected for analysis. 5 µM thick paraffin sections were dewaxed and rehydrated through graded alcohols. Heat-induced epitope retrieval was performed in a vegetable steamer using sodium citrate buffer, pH 6.0, for 25 minutes, followed by slow cooling to room temperature. Sniper background blocker (Biocare Medical, Concord CA) was applied to reduce non-specific background. For α7 nAChR, rat monoclonal Anti-Nicotinic Acetylcholine Receptor α-7 (1:20) was used (#ab24644, Abcam Inc.) followed by Alexa 647 goat anti-Rat (#A-21247, Invitrogen Inc.). For HMGB1 staining, rabbit polyclonal anti-HMGB-1 (1:250) was used (#NB100-2322, Novus Inc.) followed by Alexa 568 goat anti-rabbit (#A-11011, Invitrogen). After immunostaining, DAPI nuclear stain was used as counterstain (Molecular Probes/Invitrogen, Carslbad CA), and the section was mounted with ProLong Gold mounting medium to preserve fluorescent signal throughout the image capture (Molecular Probes/Invitrogen, Carslbad CA). All tissue sections for analysis were processed simultaneously, using pooled reagents and antibodies for consistency. Images were captured on a Zeiss AxioImager Z1 microscope, equipped with an Apotome grid-a structured illumination device, which isolates optical slices much like a confocal microscope (Carl Zeiss Canada Ltd, Toronto, ON). For each animal, eight random fields of view in each brain region were collected for analysis. All images within each brain region were captured at the same instrument settings to ensure consistent illumination and detection parameters among samples. To avoid crosstalk between channels that might create non-specific intensity signals, bandpass dichroic filters for each dye were carefully selected based on the spectral profiles of the fluorescent tags, and tested against controls that are positive for that desired dye wavelength, but negative for all other dyes used. Images were captured sequentially using one filter set at a time for each channel. The image analysis was done in Image Pro Plus 7.0 (Media Cybernetics, Bethesda, MD, USA). We had co-labeled with the HMGB1, α7 nAChR, Iba1 and DAPI, yielding 4 separate channels that we used as follows. First, we created a binary mask of the DAPI channel (white nuclei/ black background), then subtracted that from the Iba1 channel image. Since pure binary white has an intensity of 255, the subtraction left us with an Iba1 channel image with black holes of 0 intensity where the nuclei had been located, *i.e.*, an image with intensity only in the cytosolic area. Second, we created a threshold on the resulting Iba1 image to isolate the outlines of the cytosolic region only of the microglial cell bodies. Third, using the “Load Outlines” function in Image Pro, we were able to apply the outlines of the cytosolic area from the Iba1 image of the matching images, and measure data only within those outlines. Fourth, we created similar outlines of the nuclei using the DAPI channel, and applied those onto the HMGB1 and α7 nAChR channels respectively, to obtain the intensity and area data solely from the nuclear area within the outlines. All image analyses were performed on 8-bit tiff images calibrated for scale measured per pixel on a scale of 0 (black) −255 (white) within the area of interest (cytosol or nuclei). Area was calibrated to square microns. Intensity/area yielded a mean intensity/sq micron. In all analyses, samples of non-tissue background areas were measured for intensity, and positive signal histogram settings were chosen to selectively measure signal above this background. Thus, mean intensity/sq micron area of the sampling region is reported for HMGB1 and α7 nAChR. We discriminated and report separately active and quiescent microglia (aMG, qMG) based on the morphological features of the Iba1+ cells and their location in relation to the neurons: as aMG were considered Iba1+ cells exhibiting large (~2x fold) soma and processes engulfing neighbouring neurons, while qMG did not exhibit either of these features (Fig. 1). We recognize that such distinction, albeit based on general consensus regarding distinct morphological features of active (hypertrophy and increased processes) versus quiescent microglia, is also somewhat subjective, when it comes to the notion of co-localization to or engulfing of neurons. We return to this in the Discussion section.

For analyses of HMGB1 and α7 nAChR intensity per area between the brain regions within each group we had to take into account that their absolute values could not be compared due to region-specific optimized acquisition as described above. For HMGB1, we worked around this limitation by deriving relative measures of HMGB1 intensity per area as cytosol/nucleus ratio for each brain region, as HMGB1 translocation indices (the higher the ratio, the more translocation occurs), thus making them comparable.

### Fetal heart rate variability analysis to gauge activity of the cholinergic anti-inflammatory pathway

The fHRV methodology was described elsewhere. [2, 42] Briefly, R peaks were triggered by steepest ascent criterion to derive fHRV, and RMSSD was calculated from five minutes of fHRV using Matlab 6.1, R13 (The MathWorks, Natick, Massachusetts, USA). We selected ten minutes intervals of artifact-free fetal ECG for each time point. ECG-derived fHRV segments were analyzed time-matched to the cytokine samples taken at baseline and during the UCO series. In addition, fHRV measures were correlated to the brain’s white matter microglia cell counts and brain region- and cell compartment-specific α7nAChR and HMGB1 signals in the Iba1+ microglia (see above). CAP activation was measured as increases in fHRV measure RMSSD that reflects vagal modulation of fHRV. [2]

### Statistical analyses

Blood gas, pH, IL-1β and fHRV-derived measurements in response to repetitive cord occlusions were compared to the corresponding baseline values by one way repeated measures ANOVA with Holm-Sidak method of correction for multiple comparisons. A generalized estimating equations (GEE) model was used to assess the effects of UCO on HMGB1 translocation while accounting for repeated measurements in space across the brain regions with AR(1) correlation matrix. We used a linear scale response model with animal group, MG type (qMG, aMG) and brain regions as predicting factors to assess their interactions using maximum likelihood estimate and Type III analysis with Wald Chi-square statistics. A similar analysis was made to assess the behaviour of α7 nAChR intensity per area across the groups and MG type with HMGB1 translocation index as covariate, but using an independent correlation matrix (α7 nAChR intensity per area between brain regions within each group could not be compared, since absolute values had to be used; hence no repeated measurements across the brainregions were assessed for α7 nAChR intensity per area values). Correlation analysis was performed using Spearman correlation coefficient (IBM SPSS Statistics Version 21, IBM Corporation, Armonk, NY). Significance was assumed for p < 0.05. Results are provided as means ± SD or as median {25-75} percentile, as appropriate. Not all measurements were obtained for each animal studied (see Figure legends).

## Acknowledgment

The authors thank Brad Matushewski, Jac Homan, Richard Harris, Jeremy McCallum, Ashley Keen, and Maria Sinacori for technical assistance. We thank the lab of Dr. Tim Regnault who with Lin Zhao helped with establishing HMGB1 IHC in sheep. None of the authors have any competing interests in the manuscript.

